# Interrelationship of Substrate Crystallinity, Enzyme Binding Strength, and Cellulase Activity

**DOI:** 10.1101/2024.08.20.607150

**Authors:** Gustavo Avelar Molina, Kay Schaller, Jeppe Kari, Corinna Schiano-di-Cola, Günther H. J. Peters, Kim Borch, Peter Westh

**Author notes:** **Correspondence** P. Westh, Department of Biotechnology and Biomedicine, Technical University of Denmark, Søltofts Plads, building 224, DK-2800 Kongens Lyngby, Denmark.

## Abstract

Structural polysaccharides are difficult to degrade due to their crystalline structure. Hence, industrial conversion of biomass has focused on both substrate pretreatment and enzyme engineering to improve the biochemical conversion of biomass into fuels and chemicals. However, few studies have explored the interrelationship between substrate crystallinity and cellulase activity. Here, we systematically investigated the kinetics of structurally diverse cellulases on five cellulosic substrates with varying crystallinity. Regardless of enzyme structure and catalytic mechanism, we observed a linear scaling of the kinetic parameters (*K*_M_ and *k*_cat_) in a log-log plot, indicating a linear free energy relationship (LFER) between binding and activation energy. LFERs were observed for all investigated substrates, but their slopes varied distinctly and appeared linked to the substrate crystallinity. Substrates with low crystallinity exhibited LFERs with a slope near 1, while highly crystalline substrates had a slope of approximately 0.25, providing insights into the transition state (TS) for the rate-limiting step. We propose that maximal turnover was limited by slow dissociation, with the TS structurally close to the enzyme-ligand complex on crystalline substrate, while on amorphous substrate, the TS structure was closer to the dissociated system. We suggest that these observations reflect competing interactions of the ligand with respectively the enzyme binding cleft and the substrate matrix. This study emphasizes the interconnected nature of substrate pretreatment and enzyme engineering, urging a holistic approach to propel the biochemical conversion of lignocellulosic biomass, crucial for advancing sustainable production of fuels and chemicals.

## Introduction

Structural polysaccharides such as cellulose and chitin represent Nature’s largest reservoir of organic carbon and hence make attractive feedstocks for biorefineries that produce liquid fuels and chemicals [1]. The industrial process includes thermochemical pretreatment, which makes the biomass more accessible and susceptible to enzymatic hydrolysis [2]. The subsequent deconstruction into fermentable sugars (so-called saccharification) uses a cocktail of enzymes, predominantly cellulases, but the slow conversion challenges the economic feasibility of biorefineries [3–5]. For this reason, saccharification has been widely investigated, and bottlenecks linked to both enzyme- and substrate related properties have been suggested [6–8]. Substrate crystallinity has been broadly recognized as a major factor limiting the hydrolytic rate [2,9], but molecular origins of the correlation between crystallinity and enzyme efficiency remain elusive. One interpretation, which has been explored in several computational studies, is that the thermodynamic stability of the cellulose crystal determines its accessibility [10–12]. Hence, to hydrolyze the β-1-4 glucosidic bonds that link glucose moieties, cellulases need to transfer a piece of polymer chain from the substrate matrix to the enzyme’s binding cleft. For this to occur, enzyme-ligand interactions must offset the network of hydrogen bonds and hydrophobic interactions that anchor the cellulose chain to the surface. If the chain is in a conformation where such forces are weak, cellulases readily gain access to a scissile bond as little work is needed for ligand transfer [13]. For typical cellobiohydrolases (CBHs), which are the workhorses of saccharification, ligand transfer involves at least nine pyranose units, and releasing a ligand of this size from the surface of a crystalline substrate comes at a sizable free energy penalty (sometimes called the decrystallization free energy) of some 60-70 kJ/mol [12,14,15]. To overcome this penalty, CBHs form strong interactions with the cellulose strand in the Michaelis complex [16]. This balance of strong forces in both the substrate matrix and the enzyme complex is probably one of the main determinants of cellulase kinetics. It means that both binding and dissociation of the ligand may be associated with large activation barriers, and this is reflected in the observation that the rate-determining step for cellulases is binding or unbinding of the substrate, rather than the chemical steps of the hydrolytic reaction [17–20]. In addition, the turnover frequency of many cellulases is well below one per second [19], and hence much lower than typical values for hydrolases acting on soluble substrates [21], with no major penalty of substrate release. Despite the importance of the on/off dynamics of cellulolytic enzymes, the energy barriers that control these steps remain poorly explored.

We have recently investigated general kinetic trends for a wide group of almost 100 fungal cellulases on a semi-crystalline substrate (Avicel) and found that enzyme function was confined by a linear free-energy relationship (LFER) [22]. Specifically, the energy barrier of the rate-limiting step appeared to scale linearly with ligand-binding strength, even for cellulases that were structurally and mechanistically unrelated. LFER links a thermodynamic and a kinetic property, and this provides an experimental avenue to study some aspects of the energy landscape that underlies a reaction. LFERs are widely used to elucidate reaction mechanisms and substituent effects in organic chemistry [23], and they are also well-established tools within both fundamental and applied heterogeneous catalysis [24]. However, applications within biochemistry remain scarce. The most prominent examples are the seminal works by Fersht and coworkers, who used free-energy scaling within families of closely related mutants to elucidate transition states in both protein folding [25] and enzyme catalysis [26].

In the current work, we investigated the relationship between substrate crystallinity, enzyme affinity, and catalytic turnover for different cellulose-cellulase systems. Specifically, we kinetically characterized 12 cellulases belonging to five different glycosyl-hydrolases families on five different cellulosic substrates and at four temperatures. The measurements generated over 200 sets of Michaelis-Menten parameters, and this data consistently suggested linear scaling between binding and activation. The slopes of the observed LFERs differed markedly with the crystallinity of the substrate. We propose that this reflected the variation in the free energy of ligand release (ΔG_LR_) for different types of cellulose (Fig. 1) and concomitant differences in the transition state structure for the association- and dissociation steps.

**Fig. 1.**
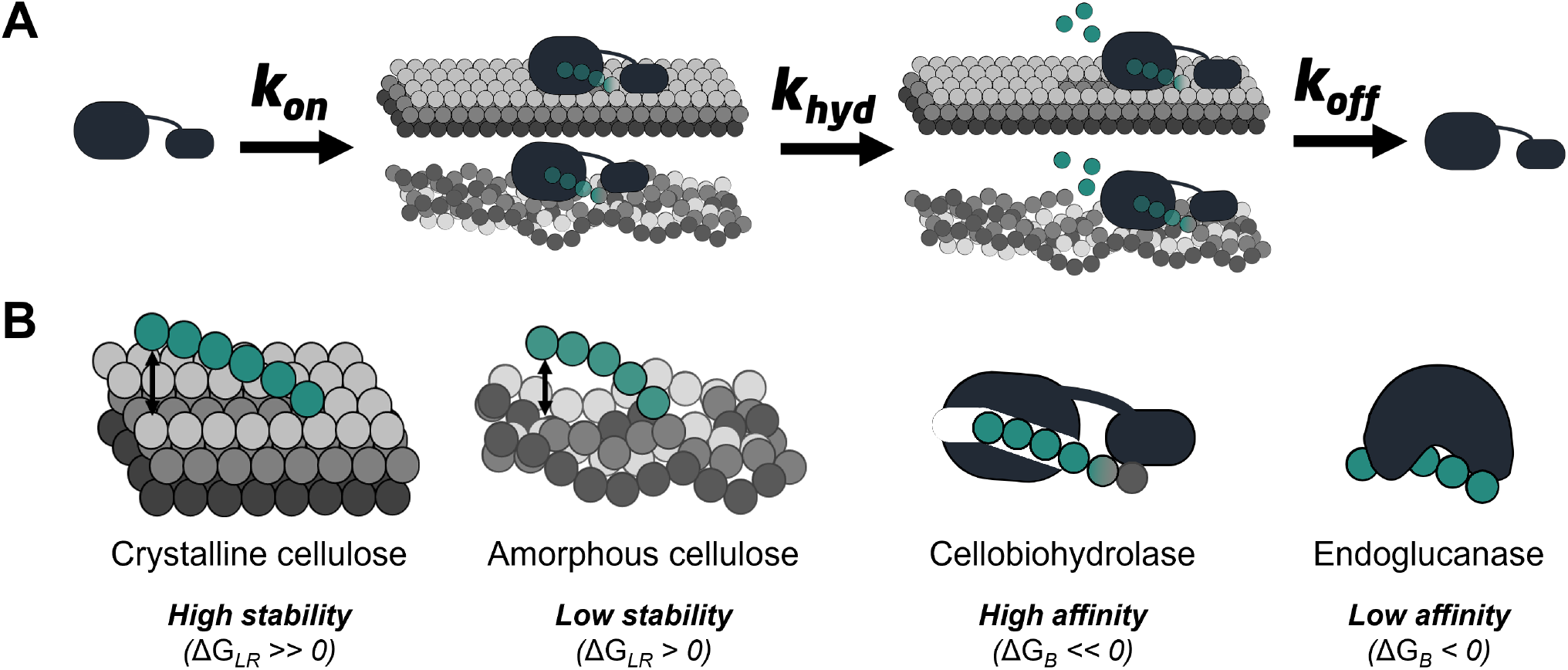
Schematic illustration of the enzymatic hydrolysis of cellulose. A) Simplified reaction scheme, which accounts for complexation, hydrolysis, and dissociation on two types of cellulose, crystalline and amorphous. B) Two types of cellulases and cellulose structures with different stability and enzyme-substrate affinity. The free energy of ligand release, ΔG_LR_, is higher for crystalline compared to amorphous cellulose. Processive cellobiohydrolases generally bind the ligand tightly (ΔG_B_ << 0), while endoglucanases have lower substrate affinity.

## Results

### Kinetic characterization of cellulases on various substrates

We kinetically characterized a group of cellulases on different substrates as detailed in Tab. 1 at four different temperatures (10 °C increments from 20 °C to 50 °C). The enzymes were selected based on two criteria. First, they covered a broad range of substrate-binding strength. Secondly, they represented several hydrolase families with different catalytic mechanisms, modes of action, and modularity. The five substrates were chosen based on their (reported) degrees of crystallinity so that high, intermediate, and low crystallinity substrates were investigated. This allowed systematic analysis of relationships between substrate crystallinity, binding affinity, and catalytic turnover.

The kinetic characterization was based on a simple Michaelis-Menten approach. Specifically, we measured catalytic rates for a constant, low enzyme concentration, at low degrees of conversion, and at variable substrate loads. Advantages and limitations of this approach for insoluble substrates (particularly cellulose) have been discussed elsewhere [27–29]. We plotted measured rates against the substrate load as exemplified in Fig. 2 and used non-linear regression with respect to the Michaelis-Menten equation (Eq. 1) to derive the kinetic parameters *K*_M_ and *k*_cat_.

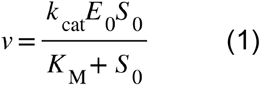

**Fig. 2.**
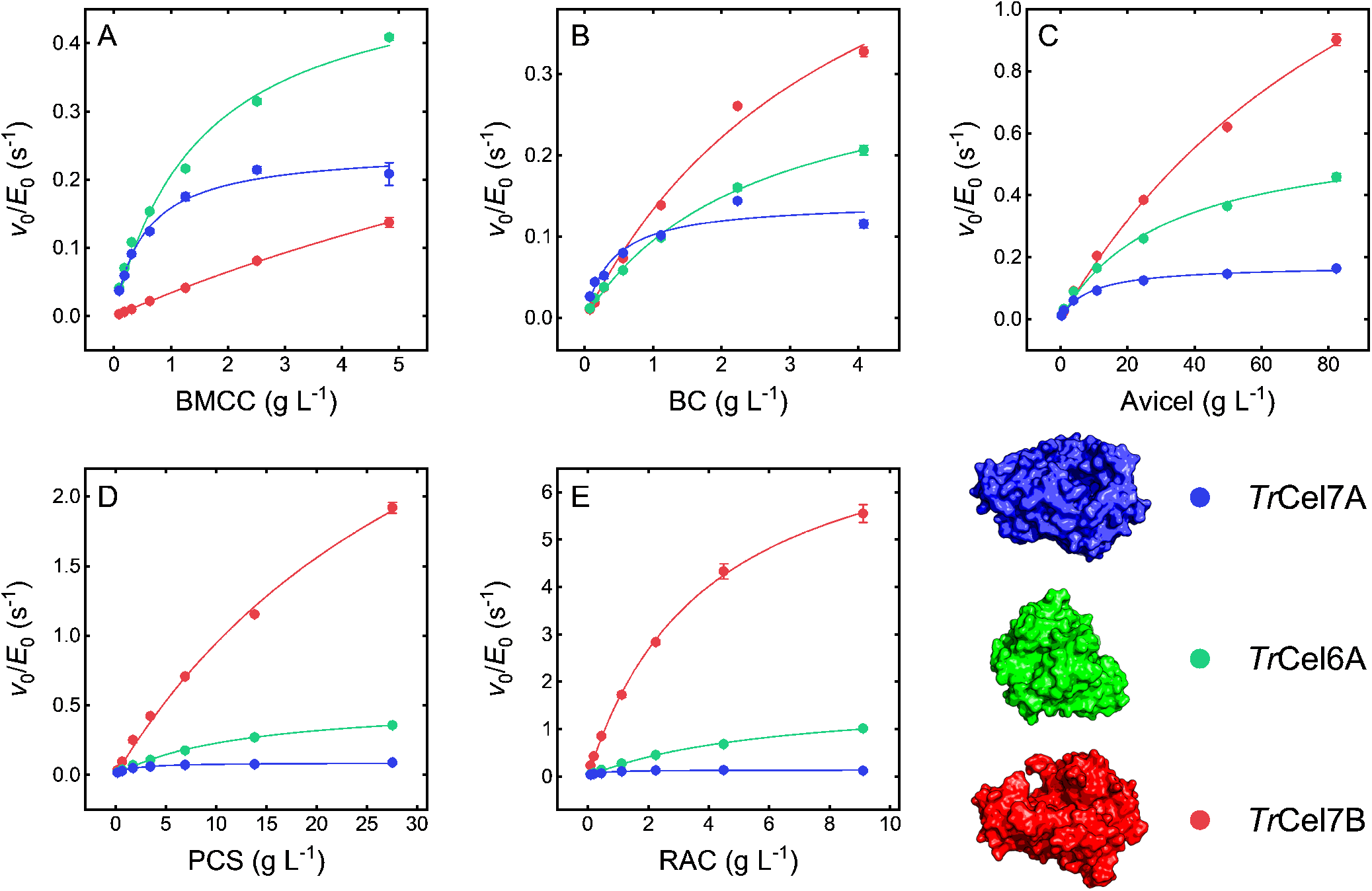
Examples of kinetic measurements and data analysis. The figure represents three enzymes from the model cellulolytic fungus *T. reesei* (*Tr*Cel7A, *Tr*Cel7B, and *Tr*Cel6B, see Tab. 1). We used five types of substrates as indicated on the abscissa of each plot (see Tab. 1 for further details regarding the substrates), and in the examples shown here, the experimental temperature was 30 °C. Symbols represent experimental data, and the lines are best fits of Eq. 1. Kinetic parameters for all combinations of enzymes, substrates, and temperatures are listed in Tab S1.

In Eq. 1, E_0_ and S_0_ are respectively the initial enzyme concentration (in M) and substrate load (in g/L). We determined the maximum likelihood values of *K*_M_ and *k*_cat_ and listed all parameters in Tab. S1 of the SI.

**Table 1.** Some key properties of the investigated cellulases and cellulosic substrates.

| Enzyme | Organism | GH | Mechanism | Modularity<br>(+/- CBM) |
| --- | --- | --- | --- | --- |
| **Cellobiohydrolases** |  |  |  |  |
| ReCel7A<sub>TrCBM</sub><sup>[a]</sup> | *Rasamsonia emersonii* | 7 | Retaining | + |
| TrCel7A | *Trichoderma reesei* | 7 | Retaining | + |
| PcCel7D | *Phanerochaete*<br>*chrysosporium* | 7 | Retaining | + |
| TrCel7A<sub>CD</sub><sup>[b]</sup> | *Trichoderma reesei* | 7 | Retaining | - |
| TrCel6A | *Trichoderma reesei* | 6 | Inverting | + |
| TrCel7A<sub>W38A</sub><sup>[c]</sup> | *Trichoderma reesei* | 7 | Retaining | + |
| TrCel6A<sub>CcCBM</sub><sup>[d]</sup> | *Trichoderma reesei* | 6 | Inverting | + |
| **Endoglucanases** |  |  |  |  |
| TrCel5A | *Trichoderma reesei* | 5 | Retaining | + |
| HiCel45A | *Humicola insolens* | 45 | Inverting | + |
| TrCel12A | *Trichoderma reesei* | 12 | Retaining | - |
| TrCel7B | *Trichoderma reesei* | 7 | Retaining | + |
| RpCel45A | *Rhizomucor pusillus* | 45 | Inverting | + |

| Substrate | Abbreviation | Crystallinity | Reference |
| --- | --- | --- | --- |
| Bacterial Microcrystalline Cellulose | BMCC | 85-90% | [19,46] |
| Bacterial Cellulose | BC | 75-85% | [61,62] |
| Microcrystalline cellulose (Avicel PH-101) | MCC/Avicel | 55-75% | [9,32,63] |
| Pretreated Corn Stover | PCS | ~50% | [63,64] |
| Regenerated Amorphous Cellulose | RAC | < 5% | [19,61] |

### Assessment of linear free-energy relationships (LFERs)

To assess possible free-energy relationships, we used ln(*k*_*cat*_) and ln(*K*_M_) as descriptors of respectively activation and binding free energy. This is a well-established approach in comparative enzymology [30,31], but we emphasize that while ln(*k*_cat_) and ln(*K*_M_) provide information on relative changes, these functions do not unambiguously reflect absolute free energies of activation and binding.

In Fig. 3, we plotted ln(*k*_*cat*_) as a function of ln(*K*_M_) for each of the five investigated cellulosic substrates (i.e., each point represents parameters for one enzyme-substrate system at one temperature). As seen from the figure, we found a linear scaling between ln(*k*_cat_) and ln(*K*_M_) for all investigated enzymes, substrates, and temperatures.

**Fig. 3.**
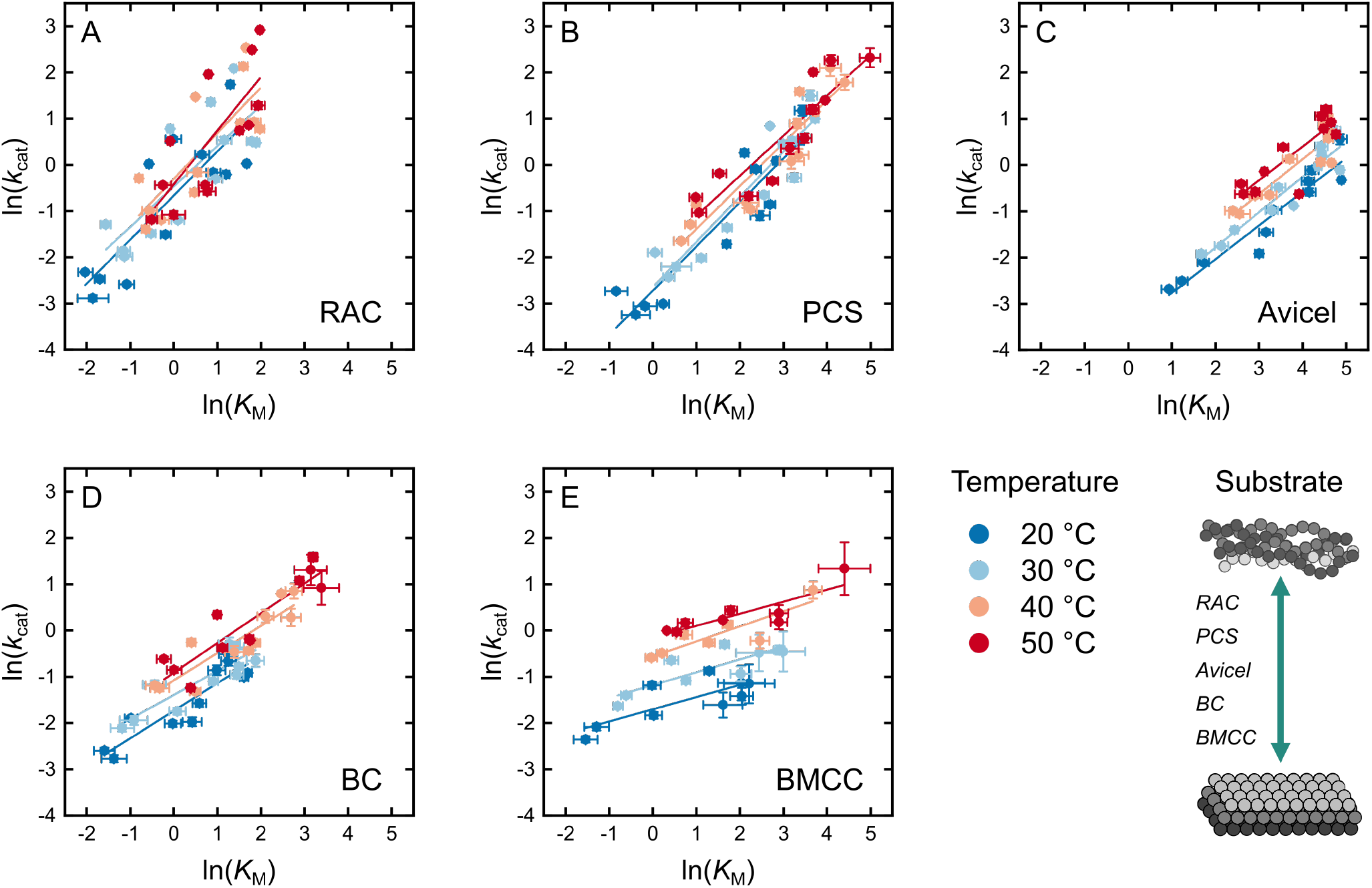
Linear free-energy relationships for cellulases at four different temperatures and five different substrates. Substrate crystallinity increases progressively from A to E. Each point represents a set of Michaelis-Menten parameters (ln(*K*_M_), ln(*k*_cat_)) for a specific enzyme, substrate, and temperature. The solid lines in all plots are derived from linear regressions to all enzymes at a fixed condition. Error bars represent standard deviations from the MM-fit (Eq. 1).

We fitted Eq. 2 to data for each substrate and temperature and listed the parameters ϕ and β together with the associated R^2^-values in Tab. S2 of the SI.

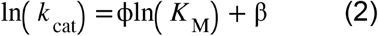

Fig. 3 consistently suggested that the cellulolytic reaction was limited by an LFER, but its location depended on temperature and substrate. In particular, the slope ϕ changed with the type of substrate. For example, the lines for RAC and PCS were steeper than lines for BMCC. Conversely, we could not detect effects of temperature on ϕ (all panels in Fig. 3 show parallel or overlapping lines).

### Calorimetric estimation of the relative crystallinity of cellulosic substrates

Fig. 3 suggested a link between substrate crystallinity and ϕ. This correlation was based on earlier measurements of crystallinity indices (CI) for related substrates (see Tab. 1), and to confirm this, we experimentally evaluated the current substrate preparations. However, absolute measurements of CI remain controversial [32,33] and beyond the current scope. Instead, we used the calorimetric method of Ioelovitch [33,34] to experimentally estimate the relative crystallinity of the current substrates. This method relies on an empirical, linear correlation between crystallinity and the enthalpy of water vapor sorption (ΔH_ads_) onto dry cellulose (ΔH_ads_ becomes less negative with increasing crystal content). Our measurements confirmed this since the measured ΔH_ads_ values scaled reasonably with CI reported previously (see SI), and we will henceforth use ΔH_ads_ as an empirical crystallinity parameter. To assess correlations between the LFERs and crystallinity, we plotted the slopes ϕ from Fig. 3 against ΔH_ads_ in Fig. 4. This graph suggested a linear decline of ϕ, and hence that LFER slopes decreased systematically as the crystallinity of the substrate increased.

**Fig. 4.**
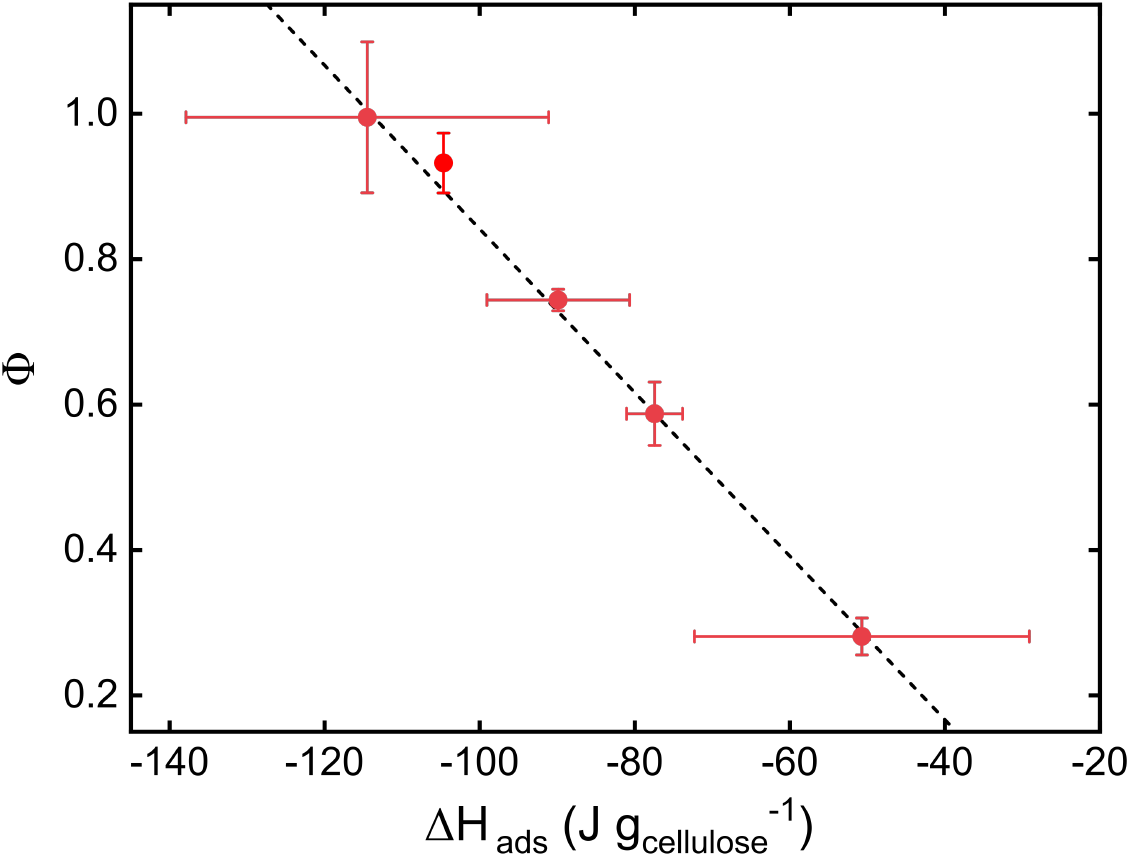
Slopes (ϕ) of the LFERs in Fig. 3 plotted as a function of the enthalpy of water vapor adsorption (ΔH_ads_) measured by isothermal calorimetry. Each data point represents one of the five investigated substrates. Earlier work [33] has shown that ΔH_ads_ provides a measure of cellulose crystallinity, and the plot implies that ϕ is about 1 for an amorphous substrate and decreases linearly with crystallinity. The dashed line derives from linear regression to the dataset.

### Computational quantification of the work to displace a cellulose strand from crystalline and amorphous cellulose matrices

To further elucidate differences between substrates, we used steered molecular dynamics simulations to study the thermodynamic penalty (see Fig. 1) of releasing a cellulose fragment with nine pyranose rings (cellononaose) from the surface of cellulose. We investigated both crystalline cellulose and an amorphous cellulose structure from an earlier computational study [35] and quantified the work of displacement, ΔG_MD_, for seven manually selected strands at four temperatures (see Methods). We enumerated atomic contacts (3.5 Å cutoff) between the cellononaose strand and cellulose prior to the displacement and normalized this number with respect to contacts in a cellononaose fragment on the edge [12] of a crystalline fibril. Hence, atomic contacts near 100% indicate that the cellononaose strand is essentially part of a crystalline structure, whereas a low percentage specifies a disordered system with sporadic ligand-surface interactions. Results in Fig. 5 showed how ΔG_MD_ increased moderately with the number of atomic contacts up to about 90% and rose steeply at higher contact numbers as the starting conformation for the MD experiment approached the crystal structure. We were not able to single out systematic effects of temperature on ΔG_MD_ in these simulations.

**Fig. 5.**
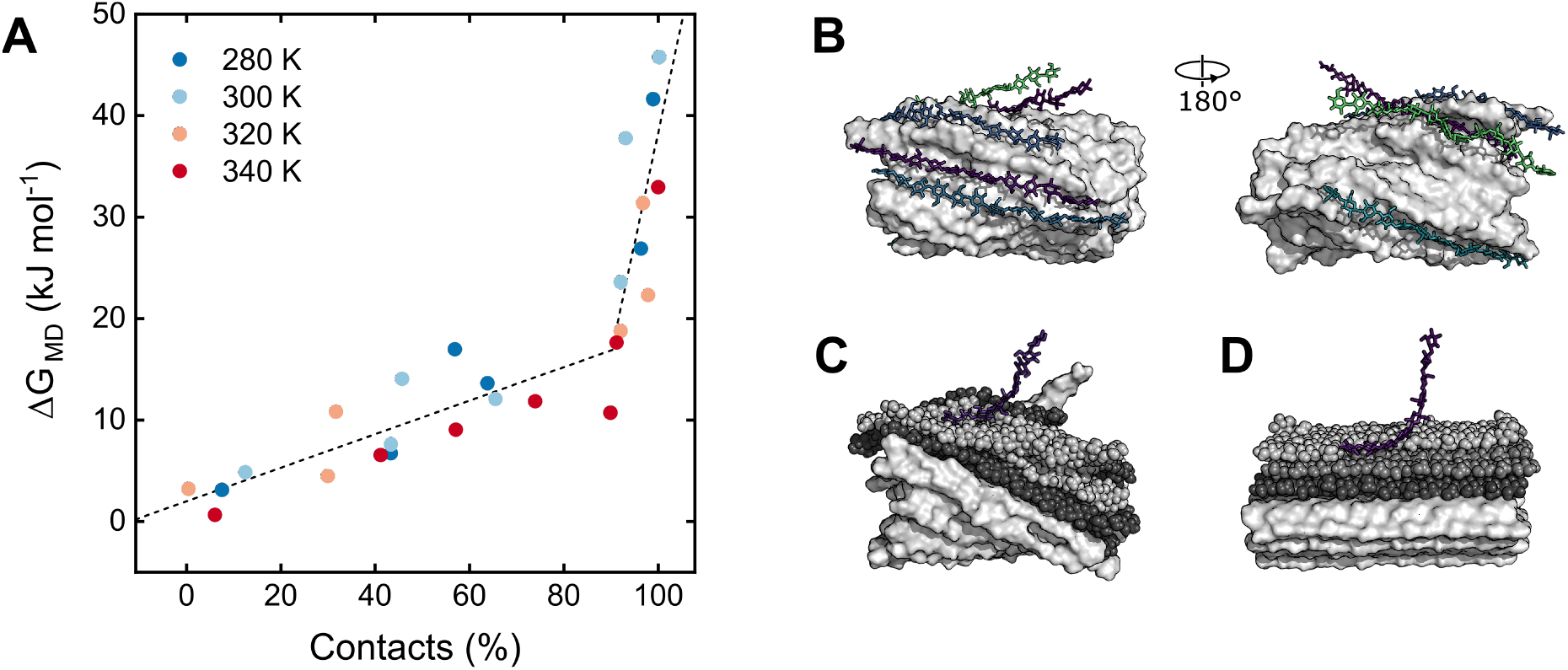
Panel A shows the free energy penalty of removing cellononaose from the surface of the cellulose matrix (ΔG_MD_) determined by steered molecular dynamics simulations. The computed free-energy change was plotted against the initial extent of ligand-surface contacts (see main text). This extent of ligand-surface contact served as a proxy of local crystal structure. Dashed lines are to facilitate visualization. The cartoons in B show examples of cellononaose ligands with different extents of surface contacts. The pictures in C and D show frames from the MD simulations with respectively amorphous and crystalline cellulose. White continuous surfaces represent remaining chains that were not part of the simulation box for context. The simulation involved the strands illustrated by spheres. Black spheres (second contact shell in the amorphous structure, third layer in the crystalline fibril) were restrained, whereas grey spheres were unrestrained.

## Discussion

The experimental data in Fig. 3 suggested scaling between Michaelis constants and the maximal turnover for a group of enzymes, which shared the same function (hydrolysis of insoluble cellulose) but varied with respect to structure and mode of action. In the following, we will discuss this observation with focus on relationships of crystallinity on enzymatic performance. Such relationships are well established empirically [9] and of direct importance for industrial applications of cellulases [2]. However, their molecular underpinnings remain to be fully understood.

The slope of an LFER can give information about the transition state (TS) of the rate-determining step in a multi-step reaction. To utilize this, we first consider the nature of rate limitation for cellulolytic enzymes. Most works on this topic have pointed towards slow dissociation as the main bottleneck, while the chemical steps of the inner hydrolytic cycle were reported to be comparably fast [17,18,20,36,37]. Using the scheme in Fig. 1, this corresponds to a reaction where *k*_hyd_ >> *k*_off_. In practice, this means that except at very low substrate loads where association inevitably becomes slow, the rate-limiting step for cellulases will be dissociation. This interpretation has been experimentally confirmed for a few cellobiohydrolases from GH6 and GH7 [17,38], and it is reflected in the schematic energy diagrams in Fig. 6, where dissociation presents the largest energy barrier in the forward direction. This directs the interest towards the structure of the TS for dissociation, and while nebulous, some experiments [39] have suggested a conformation where the ligand was partially retracted from the enzyme’s binding cleft. A free-energy apex of this structure could reflect a balance between attractive forces in the enzyme complex and the cellulose matrix (see Introduction). Hence, if the ligand moves further out of the binding cleft, it can establish better interactions with the cellulose matrix, and this would make outward movement thermodynamically downhill. Movement in the opposite direction would be favored by better enzyme-ligand interactions and hence also downhill.

**Fig. 6.**
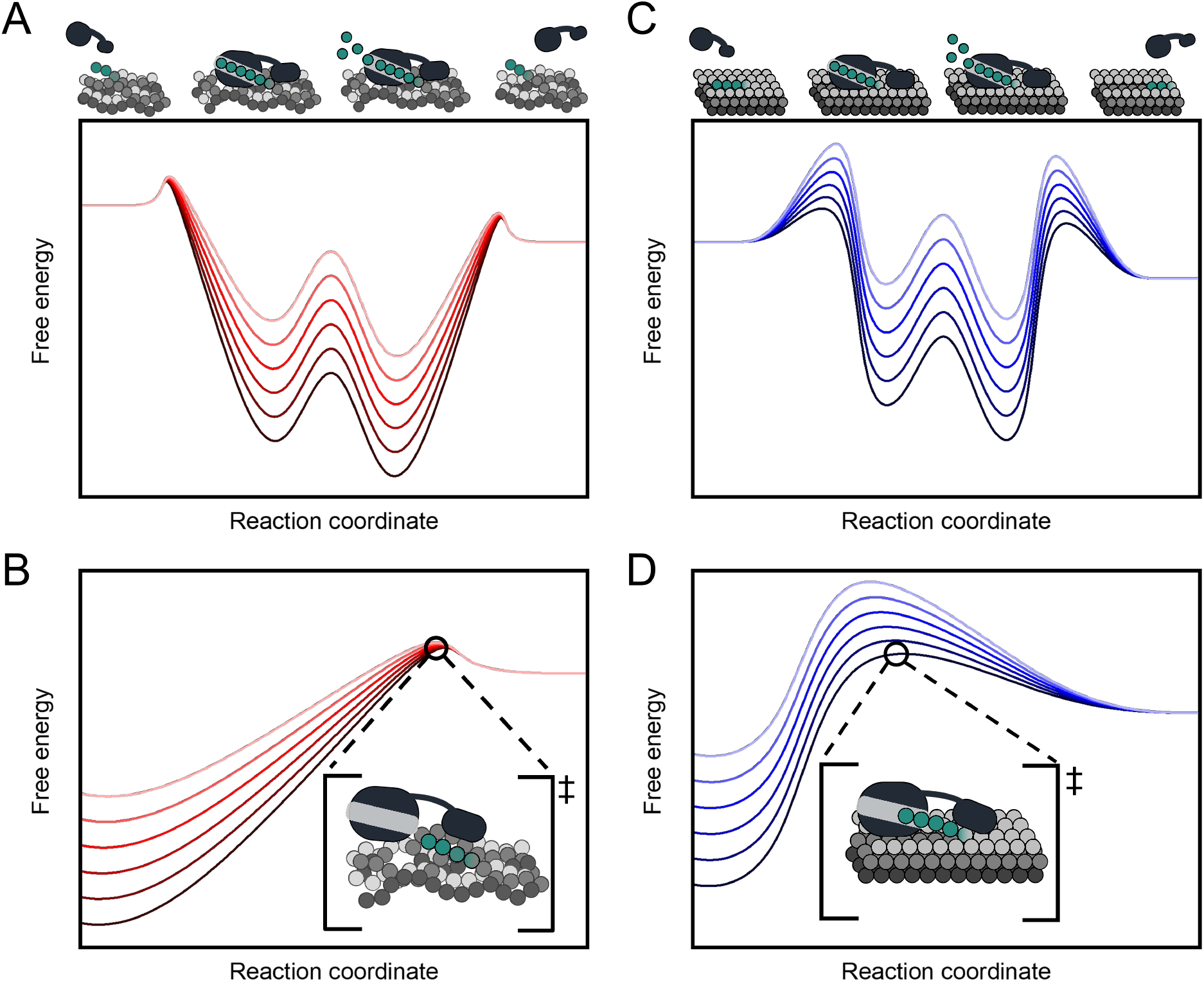
Schematic energy diagrams for the enzymatic hydrolysis of cellulose based on the simplified reaction scheme in Fig. 1. Each panel presents different enzymes attacking the same substrate. The substrate is amorphous in Panel A, and crystalline in C. Panels B and D are magnifications of the dissociation step. Curves at a low position symbolize enzymes with comparably tight substrate binding (low *K*_M_) and vice versa. For crystalline substrate (D), the free energy of the ES-intermediate and the transition state of dissociation change approximately in parallel, and this leads to a ϕ value near zero (the energy barrier of dissociation is nearly the same for different enzymes). Conversely, for amorphous cellulose (B), the different enzymes have almost the same free energy of the TS. This latter situation leads to a ϕ value near 1 (activation barrier increases commensurate with binding strength).

Returning to LFER slopes, we note that in general terms, ϕ ≈ 1 implies a TS that is structurally related to the product, whereas slopes closer to zero suggest a “reactant-like” TS structure. This relationship originates from Hammond’s postulate [23], and its meaning for cellulase dissociation is illustrated in Fig. 6. For an amorphous substrate like RAC (panel A and B), we found ϕ ≈ 1, and this means that the TS is structurally close to the dissociated system (the apex and the dissociated system are close in the reaction coordinate). Moreover, as most enzyme-ligand interactions are broken in this state, the free energy in the TS is almost the same for all enzymes on RAC (Fig. 6B). Conversely, for a crystalline substrate like BMCC (panels C and D), we found ϕ ≈ 0.25, and this means that most enzyme-ligand interactions from the ground state remain in the TS (the ES-complex and the energy apex are closer in the reaction coordinate). In other words, the TS for dissociation from BMCC is reactant-like with moderate retraction of the ligand. It is interesting that the same reaction (chemically) shows different TS structures, and we propose that this reflects the aforementioned balance between ligand interactions in enzyme-binding cleft and cellulose-surface matrix. Hence, strong ligand-matrix interactions in BMCC (Fig. 5), established as the ligand leaves the enzyme’s binding cleft, compensate for the loss of enzyme-ligand interactions. As a result, the energy apex occurs early in the dissociation path. Weaker surface binding in RAC, on the other hand, offers only poor compensation, and the TS shifts to a higher reaction coordinate with nearly full retraction of the ligand from the binding cleft.

Conceptually, LFERs of the type considered here may be expressed [40] by

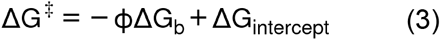

where ΔG^‡^ and ΔG_b_ are respectively the activation barrier and the standard free energy of ligand binding. For the current reaction, the intercept, ΔG_intercept_, may be understood as ΔG^‡^ in the hypothetical case where the enzyme has no net interaction with the substrate (*i*.*e*., when ΔG_b_ = 0). In this situation, the activation barrier will be entirely governed by the energetic penalty (work) of releasing the ligand from the cellulose matrix. It follows that ΔG_intercept_ is related to this work. Under this interpretation, Eq. 3 singles out enzyme-related (ΔG_b_) and substrate-related (ΔG_intercept_ and ϕ) contributions to the overall activation barrier. To test this view against the experimental data, we compared crystallinity and the descriptor for the activation barrier used here, ln(*k*_cat_). Specifically, we plotted -ln(*k*_cat_) as a function of the calorimetric crystallinity parameter, ΔH_ads_, for the three *T. reesei* enzymes (the same enzymes as in Fig. 2).

Results in Fig. 7 may be rationalized with respect to Eq. 3. For amorphous cellulose, for example, ligand release from the surface is energetically inexpensive (Fig. 5). As argued above, this corresponds to a small value of ΔG_intercept_, and the activation barrier may be approximated ΔG^‡^ ∼ –ϕΔG_b_. This is in accord with Fig. 7, which shows that the activation barrier on amorphous substrate is strongly coupled to ΔG_b_. Specifically, Cel7A, with tight ligand binding (large negative ΔG_b_), has a much higher barrier than a weak binder like Cel7B, and we conclude that on amorphous substrates, the activation barrier is predominantly controlled by an enzyme property – ligand binding strength. For crystalline substrates, on the other hand, ligand release is costly (large ΔG_intercept_) and ϕ is small (Figs. 4 and 5), and as a result, Eq. 3 may be approximated to ΔG^‡^ ∼ ΔG_intercept_. This latter approximation suggests that the activation barrier is almost independent of ligand binding strength. This is also in line with Fig. 7, as the different enzymes had approximately the same activation barrier on crystalline substrates. We deduce that the activation barrier on a crystalline substrate is predominantly governed by a substrate property - ligand release. Overall, Fig. 7 shows that the activation barrier scales with crystallinity between the two limiting cases discussed above. For Cel7A, with tight ligand binding, the slope was negative, whereas the weak binder Cel7B had a positive slope. This probably reflected dominance of the linear term (–ϕΔG_b_) and the intercept (ΔG_intercept_) in Eq. 3 for respectively strong- and weak binders. For an enzyme with intermediate binding strength (Cel6A), these effects appeared to compensate each other, and consequently, the activation barrier showed little dependence on crystallinity. As a final aspect of Fig. 7, we note that the minimal activation barriers observed for Cel7A and Cel7B at respectively high and low crystallinity parallel the substrate specificity typically reported for these enzymes [3].

**Fig. 7.**
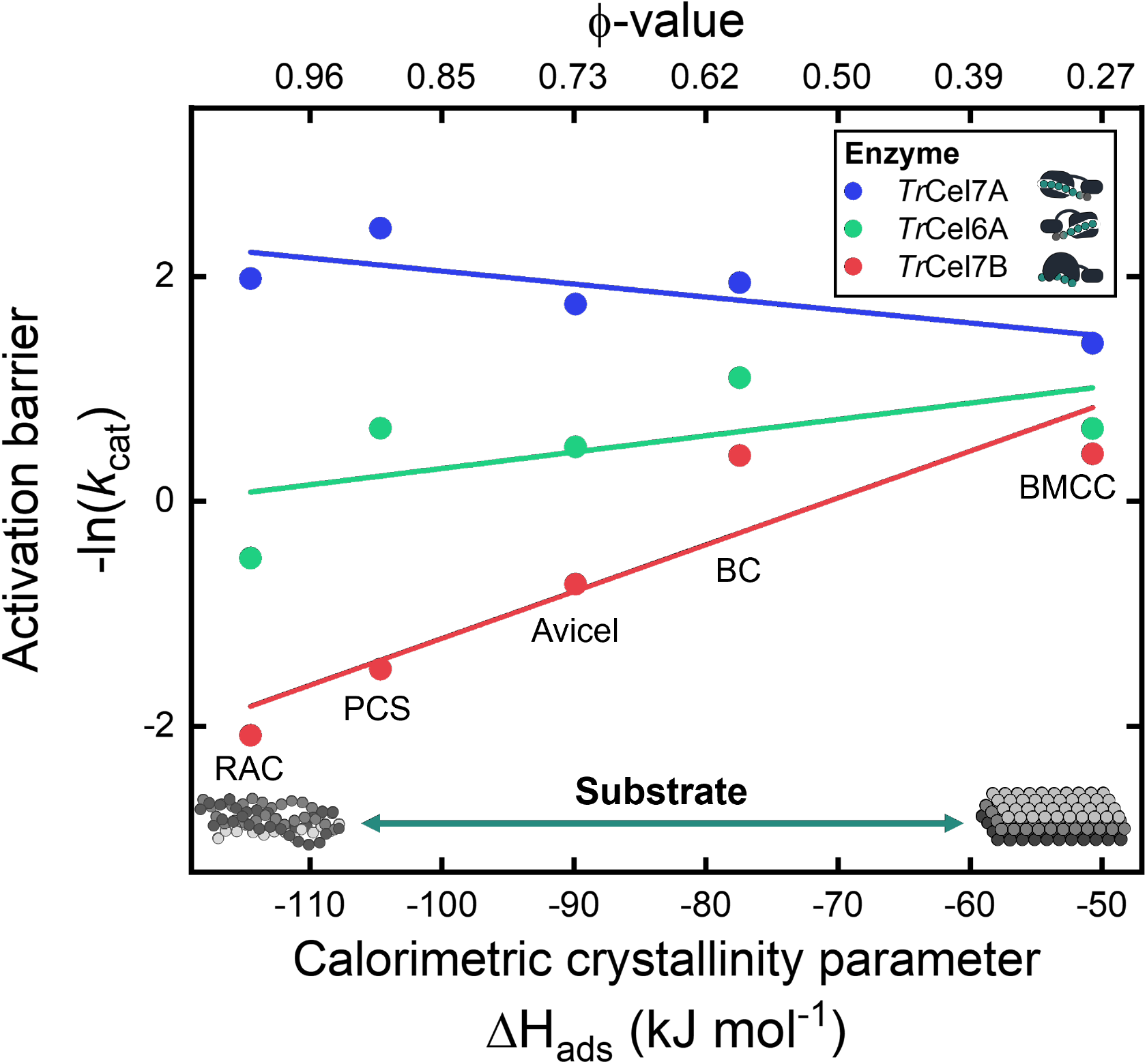
Activation barrier for the rate-limiting step (expressed as -ln(*k*_cat_)) for three *T. reesei* enzymes acting on different substrates. The lower abscissa is the crystallinity parameter derived from calorimetric measurements (see Fig. S3), and on the upper axis, this is converted into to slope of the LFERs, ϕ, according to Fig. 4. Solid lines derive from linear regression to the datasets of corresponding color.

It is known that biomass pretreatment makes the recalcitrant substrate more amenable to attack by cellulases [3]. This process lowers the penalty of ligand release (Fig. 5) and hence promotes substrate accessibility (see Introduction). However, these benefits may be linked to an enlargement of the activation barrier of dissociation (Fig. 7A), and the overall effect consequently depends on ligand-binding strength for the applied enzymes as well as the substrate load [41]. Similar considerations can be used for comparisons of different enzymes (wild types or engineered variants). A CBH engineered to high substrate affinity, for example, excels with respect to attacks on tightly anchored ligands on a crystalline fibril. However, as illustrated in Figs. 6 and 7, this advantage may be counteracted by a very low *k*_cat_ on more disordered substrates, and high substrate affinity may hence be a drawback on thoroughly pretreated biomass.

In closing, we note that enzymatic hydrolysis of cellulose and other insoluble, polymeric substrates has the unusual trait that retrieval of the ligand requires considerable work. This creates a particular need for strong, compensating enzyme-ligand interactions, which appear to be met by the evolution of long clefts with tight substrate binding in cellulases. The current work provided some hints about how the balance of forces between the substrate matrix and the enzyme complex collectively govern cellulase function. When comparing different cellulases that attack an amorphous substrate, we found that binding and activation free energies for respectively complexation and dissociation changed in parallel (Fig. 6A and 6B). As a result, tight enzyme-substrate binding was associated with very slow turnover. For crystalline substrates, on the other hand, this coupling was much weaker, and consequently, the activation barrier for dissociation was only weakly affected by the strength of enzyme-ligand interactions (Fig. 6C and 6D). We propose that these general relationships between interaction strength and enzyme function can be used as a qualitative framework to rationalize cellulase performance. It may, for example, help elucidating the effects of enzyme engineering or different pretreatment protocols, which are known to alter cellulose crystallinity. Finally, this dual effect of cellulase-cellulose interaction suggests that an advanced biorefinery industry will require the concomitant and combined development of both enzyme engineering and biomass pretreatment protocols in order to enhance cellulose conversion efficiency.

## Materials and Methods

### Enzyme production and characterization

We used 12 cellulases produced by heterologous expression in *Aspergillus oryzae* and purified as described previously [22,42,43]. These earlier works also describe the methods used for the determination of enzyme purity and concentration. The activity of purified enzymes on different types and loads of cellulose was measured by the para-hydroxybenzoic acid hydrazide method [44], as detailed elsewhere [29,42].

### Substrate preparation

We investigated five types of cellulose. Avicel PH-101 was a commercial product from Fluka, and it was used as delivered. Regenerated amorphous cellulose (RAC) was produced by phosphorous acid swelling of Avicel following the procedure of Zhang and coworkers [45]. We extracted cellulose produced by the bacterium *Gluconoacetobacter xylinus* from commercial nata de coco (coconut gel in syrup, Chaokoh, Thailand) as described elsewhere [19], and we will henceforth refer to this substrate as Bacterial Cellulose (BC). A fraction of the BC preparation was further processed to obtain the so-called Bacterial Microcrystalline Cellulose (BMCC) using the method by Väljamäe et al [46]. The last substrate, Pretreated Corn Stover (PCS), was produced by the National Renewable Energy Laboratory (NREL) [47] and prepared for activity measurements as described earlier [48]. A list of all investigated enzymes and substrates can be found in Tab. 1.

### Calorimetric crystallinity determination

Substrate crystallinity was assessed from measurements of the enthalpy of water vapor sorption to dried cellulose samples. This method rests on the observation of a linear relationship between the crystallinity of a cellulose sample and the exothermic heat of its wetting [33,34]. We used a thermal activity monitor (TAM) 2277 (Thermometric A/B, Järfälla, Sweden) equipped with a 2250 RH perfusion ampoule (Thermometric) for controlled relative humidity and a purge gas. Cellulose for calorimetric measurements was initially washed at least five times with deionized water by centrifugation and resuspension. Aqueous suspensions with approximately 10 mg dry weight of cellulose were transferred into the calorimetric vessel (1 ml steel cell), flash-frozen in liquid nitrogen, thoroughly lyophilized, and stored in a desiccator. Before use, the cellulose samples were dried again at 90°C for 5-10 min, weighed, and transferred to the calorimeter. The calorimetric cell was perfused by a nitrogen gas flow (100 ml/h), which was dry (relative humidity, RH = 0%) for the first 3 h. Subsequently, the RH of the perfusion gas was ramped linearly from 0% to 96% over 5 h and kept constant at 96% for an additional 15 h to obtain equilibrium. The measured heat flow as a function of time was used to calculate the total heat of hydration for each cellulosic sample. The experimental temperature was 25 °C. The calorimetric cells were weighed immediately after hydration to obtain the mass of adsorbed water.

### MD simulation

*Simulations of the Crystalline Substrate*. A crystalline cellulose fibril with a length of 13 sugar units was set up using the Cellulose Builder webserver [49]. An edge chain (see Fig. S4B) was taken and split into a nonaose. The CHARMM36 force field was used to describe the system [50–53]. All simulations were run in GROMACS 2018.6 [54–57]. GROMACS was used to construct a cuboid box with distances of 1.2 nm towards the x- and z-faces of the box and 8.2 nm, ensuring enough space to pull the nonaose in y-direction. The systems were solvated with TIP3P water [58]. Minimization was conducted in a steepest-descent over 10’000 iterations. All subsequent simulations were performed at 300 K. An NVT-simulation over 100 ps and an NPT-simulation over 100 ps were done in succession, while keeping the whole crystal restraint. For all further simulation, only the lower-most layer was position restraint. Afterwards, an NPT-simulation over 1 ns was performed to fully equilibrate the system. Then a steered simulation over 640 ps was run, while pulling the nonaose from the crystal with a pulling rate of 0.01 nm/ps and a fore constant of 1000 kJ/mol/nm^2^. The resulting trajectories were used to prepare further simulations. Frames every travelled 0.5 Å by the ligand were extracted up to a final distance of 1.5 nm between the nonaose and the crystal. The extracted frames were used as starting configuration for Umbrella sampling simulation along the binding path (see Fig. S5 for a scheme of the method) [59]. Each window was again equilibrated over 100 ps and then a simulation for each was run 5 ns with umbrella sampling while keeping the distance restraint.

#### Simulation of the Amorphous Substrate

The structure of Vermaas *et. al* [35] was taken and a single amorphous fibril was extracted. The fibril length was reduced to the first 13 sugar units. To generate simulation with a different amount of amorphousness, several chains (C0, C2, C3, O16, O3, O4; see Fig. S4A) from the fibrils surface were prepared. That chain of interest was shortened to a nonaose and a first shell of all chains with atoms within 3 Å around the selected chain was selected, followed by a second shell of all chains with atoms within 3 Å distance around the first shell (see Fig. S4C). These structures were used and the same workflow as for the crystalline substrate was followed. For the NVT- and the first NPT-simulation, all the chains were restraint. For later simulation only the chains in the second shell were restrained.

#### Analysis

Analysis of the trajectories was performed with GROMACS. The weighted histogram analysis method (WHAM [60]) was applied to analyze the Umbrella sampling simulations along the binding path [59]. If density gaps occurred, additional windows at those distances were inserted iteratively until no gaps occurred. From the resulting potential of mean force (PMF) curves, the energy difference between minima and maxima of the curves were calculated. The errors were estimated with bootstrapping. The number of contacts between the nonaose and the other crystal chains was calculated in the 5 ns simulation of the first window (unpulled chain) for the crystalline and all amorphous simulations. The fraction of established contacts was used as a proxy to describe the amount of local crystallinity for each of the amorphous chains.

## Author contributions

GAM, JK, and PW conceptualized the manuscript and the experimental methodology. GAM planned and performed all experiments and analyzed the data. KS conceptualized the computational methodology. KS produced and analyzed the computational data. GAM, JK, and PW performed the theoretical interpretation of all results. GAM, KS, JK, and PW wrote the manuscript. PW and KB contributed with funding acquisition. KB and CSdC provided general insights into the project. GHJP provided insights into the computational method and analysis. JK, PW, CSdC, KB, and GHJP contributed with supervision.

## Acknowledgements

The work was conducted as part of the Sabatier Project, a collaboration between the Technical University of Denmark and Novozymes A/S funded mainly by Independent Research Fund Denmark [Grant number: 8022-00165B] and Novo Nordisk Foundation [Grant number: NNF15OC0016606 and NNF17SA0028392]. In addition, funding was provided by Innovation Fund Denmark [Grant number: 5150-00020B].

## Abbreviations

CBH: cellobiohydrolase
LFER: linear free-energy relationship
BMCC: bacterial microcrystalline cellulose
BC: bacterial cellulose
PCS: pretreated corn stover
RAC: regenerated amorphous cellulose
GH: glycoside hydrolase
CBM: carbohydrate-binding module
MM: Michaelis-Menten
CI: crystallinity index
MD: molecular dynamics
TS: transition-state
ES: enzyme-substrate.

