## Supplementary information for "Interrelationship of Substrate Crystallinity, Enzyme Binding Strength, and Cellulase Activity"

##### This PDF includes:

Figs. S1 to S5

Tables S1 and S2

SI references

### Supplementary text

#### Cellulase characterization parameters

**Table S1.** Kinetic parameters for 12 cellulases characterized on BMCC, BC, Avicel, PCS, and RAC at 4 different temperatures.

| Enzyme | 20 °C | | 30 °C | | 40 °C | | 50 °C | |
|---|---|---|---|---|---|---|---|---|
| | $k_{cat}$ ($s^{-1}$) | $K_M$ ($g\ L^{-1}$) | $k_{cat}$ ($s^{-1}$) | $K_M$ ($g\ L^{-1}$) | $k_{cat}$ ($s^{-1}$) | $K_M$ ($g\ L^{-1}$) | $k_{cat}$ ($s^{-1}$) | $K_M$ ($g\ L^{-1}$) |
| **BMCC** | | | | | | | | |
| *Re*Cel7A$_{TrCBM}$ | 0.095 ± 0.006 | 0.21 ± 0.06 | 0.194 ± 0.006 | 0.45 ± 0.05 | 0.552 ± 0.027 | 0.97 ± 0.13 | 0.993 ± 0.016 | 1.38 ± 0.05 |
| *Tr*Cel7A | 0.124 ± 0.009 | 0.28 ± 0.08 | 0.245 ± 0.010 | 0.54 ± 0.07 | 0.610 ± 0.029 | 1.23 ± 0.15 | 0.965 ± 0.038 | 1.73 ± 0.16 |
| *Pc*Cel7D | 0.159 ± 0.011 | 1.02 ± 0.20 | 0.337 ± 0.016 | 2.14 ± 0.21 | 0.765 ± 0.055 | 3.64 ± 0.48 | 1.255 ± 0.030 | 4.96 ± 0.19 |
| *Tr*Cel7A$_{CD}$ | 0.320 ± 0.119 | 7.66 ± 4.16 | 0.615 ± 0.245 | 11.51 ± 6.05 | N/A | N/A | N/A | N/A |
| *Tr*Cel6A | 0.305 ± 0.022 | 0.98 ± 0.20 | 0.523 ± 0.035 | 1.53 ± 0.25 | 0.906 ± 0.077 | 2.06 ± 0.38 | 1.175 ± 0.095 | 2.10 ± 0.37 |
| *Tr*Cel7A$_{W38A}$ | 0.199 ± 0.055 | 5.01 ± 2.26 | 0.391 ± 0.064 | 7.57 ± 1.82 | 0.800 ± 0.132 | 11.68 ± 2.55 | 1.450 ± 0.253 | 18.08 ± 3.82 |
| *Tr*Cel6A$_{CcCBM}$ | 0.418 ± 0.026 | 3.65 ± 0.41 | 0.740 ± 0.049 | 5.20 ± 0.56 | 1.140 ± 0.088 | 5.57 ± 0.68 | 1.544 ± 0.140 | 6.03 ± 0.85 |
| *Tr*Cel5A | 0.241 ± 0.043 | 7.70 ± 2.01 | 0.632 ± 0.271 | 19.91 ± 10.18 | N/A | N/A | 1.198 ± 0.188 | 18.07 ± 3.43 |
| *Hi*Cel45A | N/A | N/A | N/A | N/A | N/A | N/A | N/A | N/A |
| *Tr*Cel12A | N/A | N/A | N/A | N/A | N/A | N/A | N/A | N/A |
| *Tr*Cel7B | 0.316 ± 0.133 | 9.14 ± 5.36 | 0.655 ± 0.066 | 18.18 ± 2.20 | 2.391 ± 0.442 | 39.79 ± 8.10 | 3.796 ± 2.159 | 81.33 ± 48.57 |
| *Rp*Cel45A | N/A | N/A | N/A | N/A | N/A | N/A | N/A | N/A |
| **BC** | | | | | | | | |
| *Re*Cel7A$_{TrCBM}$ | 0.063 ± 0.005 | 0.25 ± 0.07 | 0.121 ± 0.009 | 0.31 ± 0.08 | 0.304 ± 0.021 | 0.64 ± 0.13 | 0.536 ± 0.030 | 0.79 ± 0.13 |
| *Tr*Cel7A | 0.074 ± 0.004 | 0.20 ± 0.05 | 0.143 ± 0.013 | 0.40 ± 0.13 | 0.287 ± 0.021 | 0.72 ± 0.16 | 0.426 ± 0.028 | 1.01 ± 0.17 |
| *Pc*Cel7D | 0.150 ± 0.006 | 0.37 ± 0.05 | 0.311 ± 0.027 | 0.64 ± 0.17 | 0.769 ± 0.031 | 1.50 ± 0.14 | 1.402 ± 0.060 | 2.71 ± 0.22 |
| *Tr*Cel7A$_{CD}$ | 0.138 ± 0.013 | 1.52 ± 0.34 | 0.385 ± 0.035 | 4.24 ± 0.64 | 0.644 ± 0.041 | 5.57 ± 0.53 | 0.811 ± 0.022 | 5.73 ± 0.23 |
| *Tr*Cel6A | 0.206 ± 0.015 | 1.80 ± 0.28 | 0.333 ± 0.018 | 2.50 ± 0.26 | 0.629 ± 0.082 | 4.02 ± 0.88 | 0.688 ± 0.043 | 3.04 ± 0.35 |
| *Tr*Cel7A$_{W38A}$ | 0.133 ± 0.009 | 0.98 ± 0.18 | 0.174 ± 0.013 | 1.09 ± 0.21 | 0.267 ± 0.015 | 1.63 ± 0.20 | 0.289 ± 0.011 | 1.46 ± 0.12 |
| *Tr*Cel6A$_{CcCBM}$ | 0.399 ± 0.026 | 5.45 ± 0.54 | 0.458 ± 0.026 | 4.46 ± 0.41 | 0.763 ± 0.057 | 6.50 ± 0.71 | 0.834 ± 0.052 | 5.64 ± 0.53 |
| *Tr*Cel5A | 0.596 ± 0.072 | 3.47 ± 0.73 | 0.744 ± 0.106 | 3.57 ± 0.88 | 2.215 ± 0.053 | 11.79 ± 0.35 | 4.887 ± 0.446 | 24.42 ± 2.53 |
| *Hi*Cel45A | 0.423 ± 0.044 | 2.68 ± 0.53 | 0.694 ± 0.080 | 4.16 ± 0.79 | 2.341 ± 0.381 | 15.75 ± 3.09 | 3.693 ± 1.208 | 23.17 ± 8.66 |
| *Tr*Cel12A* | 0.979 ± 0.149 | 19.39 ± 3.46 | 1.288 ± 0.475 | 21.77 ± 9.24 | N/A | N/A | N/A | N/A |
| *Tr*Cel7B | 0.516 ± 0.050 | 3.50 ± 0.60 | 0.666 ± 0.092 | 3.99 ± 0.92 | 1.356 ± 0.188 | 8.19 ± 1.56 | 2.932 ± 0.239 | 17.84 ± 1.72 |
| *Rp*Cel45A | 0.362 ± 0.022 | 5.02 ± 0.48 | 0.519 ± 0.071 | 6.53 ± 1.30 | 1.322 ± 0.245 | 14.75 ± 3.33 | 2.500 ± 0.914 | 29.51 ± 12.00 |
| **Avicel** | | | | | | | | |
| *Re*Cel7A$_{TrCBM}$ | 0.068 ± 0.002 | 2.6 ± 0.4 | 0.146 ± 0.004 | 5.3 ± 0.6 | 0.371 ± 0.019 | 11.0 ± 2.0 | 0.665 ± 0.026 | 13.3 ± 1.7 |
| *Tr*Cel7A | 0.082 ± 0.002 | 3.4 ± 0.4 | 0.173 ± 0.007 | 8.5 ± 1.3 | 0.346 ± 0.025 | 12.9 ± 3.0 | 0.533 ± 0.033 | 14.1 ± 2.8 |
| *Pc*Cel7D | 0.121 ± 0.004 | 5.6 ± 0.7 | 0.246 ± 0.006 | 11.4 ± 0.9 | 0.513 ± 0.023 | 18.4 ± 2.4 | 0.869 ± 0.022 | 22.7 ± 1.5 |
| *Tr*Cel7A$_{CD}$ | 0.147 ± 0.005 | 20.2 ± 1.9 | 0.412 ± 0.004 | 44.6 ± 0.8 | 1.069 ± 0.058 | 82.7 ± 7.6 | 1.946 ± 0.109 | 116.7 ± 9.9 |
| *Tr*Cel6A | 0.377 ± 0.027 | 28.1 ± 5.0 | 0.616 ± 0.040 | 31.4 ± 4.9 | 1.137 ± 0.088 | 40.7 ± 6.8 | 1.471 ± 0.077 | 34.9 ± 4.2 |
| *Tr*Cel7A$_{W38A}$ | 0.234 ± 0.014 | 23.6 ± 3.7 | 0.375 ± 0.021 | 26.8 ± 3.9 | 0.524 ± 0.033 | 25.4 ± 4.1 | 0.561 ± 0.018 | 18.5 ± 1.7 |
| *Tr*Cel6A$_{CcCBM}$ | 0.558 ± 0.044 | 63.6 ± 9.3 | 0.970 ± 0.043 | 78.1 ± 6.0 | 1.792 ± 0.111 | 98.7 ± 9.8 | 2.187 ± 0.090 | 88.2 ± 6.0 |
| *Tr*Cel5A | 0.896 ± 0.077 | 67.6 ± 10.5 | 1.500 ± 0.127 | 84.1 ± 12.0 | 2.339 ± 0.191 | 90.6 ± 12.1 | 2.882 ± 0.238 | 83.6 ± 11.6 |
| *Hi*Cel45A | 0.703 ± 0.056 | 61.9 ± 9.2 | 1.229 ± 0.069 | 84.7 ± 8.0 | 2.008 ± 0.134 | 104.9 ± 10.9 | 2.502 ± 0.125 | 104.1 ± 8.1 |
| *Tr*Cel12A | 0.723 ± 0.030 | 132.5 ± 8.1 | 0.898 ± 0.044 | 129.2 ± 9.3 | 1.044 ± 0.041 | 105.8 ± 6.5 | 0.533 ± 0.022 | 50.1 ± 4.1 |
| *Tr*Cel7B | 1.747 ± 0.190 | 129.0 ± 20.7 | 2.088 ± 0.213 | 111.2 ± 17.4 | 2.768 ± 0.204 | 96.6 ± 11.4 | 3.340 ± 0.241 | 93.4 ± 10.9 |
| *Rp*Cel45A# | 0.678 ± 0.178 | 168.4 ± 60.8 | 0.429 ± 0.037 | 70.3 ± 10.8 | 0.194 ± 0.017 | 14.6 ± 4.0 | 0.105 ± 0.009 | 7.3 ± 2.6 |
| **PCS** | | | | | | | | |
| *Re*Cel7A$_{TrCBM}$ | 0.047 ± 0.003 | 0.84 ± 0.22 | 0.111 ± 0.010 | 1.72 ± 0.59 | 0.274 ± 0.009 | 2.37 ± 0.29 | 0.496 ± 0.021 | 2.68 ± 0.40 |
| *Tr*Cel7A | 0.039 ± 0.002 | 0.68 ± 0.22 | 0.088 ± 0.003 | 1.44 ± 0.19 | 0.192 ± 0.008 | 1.92 ± 0.31 | 0.355 ± 0.017 | 2.89 ± 0.48 |
| *Pc*Cel7D | 0.065 ± 0.003 | 0.43 ± 0.11 | 0.150 ± 0.005 | 1.05 ± 0.16 | 0.424 ± 0.012 | 2.71 ± 0.27 | 0.833 ± 0.043 | 4.63 ± 0.71 |
| *Tr*Cel7A$_{CD}$ | 0.049 ± 0.002 | 1.27 ± 0.18 | 0.133 ± 0.005 | 3.05 ± 0.35 | 0.380 ± 0.023 | 9.47 ± 1.36 | 0.707 ± 0.044 | 15.61 ± 1.92 |
| *Tr*Cel6A | 0.334 ± 0.033 | 11.72 ± 2.58 | 0.522 ± 0.028 | 12.90 ± 1.49 | 1.077 ± 0.175 | 24.19 ± 6.80 | 1.427 ± 0.167 | 23.19 ± 4.77 |
| *Tr*Cel7A$_{W38A}$ | 0.180 ± 0.006 | 5.44 ± 0.49 | 0.256 ± 0.010 | 5.52 ± 0.61 | 0.441 ± 0.053 | 8.55 ± 2.50 | 0.510 ± 0.043 | 9.02 ± 1.83 |
| *Tr*Cel6A$_{CcCBM}$ | 0.423 ± 0.025 | 14.81 ± 1.79 | 0.758 ± 0.070 | 25.77 ± 4.04 | 1.233 ± 0.168 | 28.53 ± 6.39 | 1.786 ± 0.175 | 32.70 ± 5.06 |
| *Tr*Cel5A | 1.295 ± 0.045 | 8.22 ± 0.70 | 2.308 ± 0.033 | 14.78 ± 0.43 | 4.869 ± 0.279 | 29.03 ± 2.73 | 7.467 ± 0.333 | 39.82 ± 2.66 |
| *Hi*Cel45A | 0.910 ± 0.049 | 10.67 ± 1.30 | 1.567 ± 0.090 | 19.98 ± 2.12 | 2.443 ± 0.250 | 27.57 ± 4.68 | 3.320 ± 0.309 | 38.99 ± 5.46 |
| *Tr*Cel12A | 1.087 ± 0.044 | 17.09 ± 1.34 | 1.682 ± 0.094 | 24.59 ± 2.36 | 3.172 ± 0.136 | 41.79 ± 2.64 | 4.052 ± 0.130 | 52.54 ± 2.34 |
| *Tr*Cel7B | 3.235 ± 0.349 | 30.93 ± 5.36 | 4.448 ± 0.480 | 36.99 ± 6.09 | 8.177 ± 1.482 | 58.90 ± 14.45 | 9.634 ± 1.064 | 60.19 ± 8.95 |
| *Rp*Cel45A | 1.672 ± 0.129 | 28.29 ± 3.60 | 2.707 ± 0.150 | 41.46 ± 3.40 | 5.952 ± 0.934 | 81.90 ± 16.24 | 10.186 ± 2.097 | 146.47 ± 34.78 |
| **RAC** | | | | | | | | |
| *Re*Cel7A$_{TrCBM}$ | 0.075 ± 0.003 | 0.34 ± 0.06 | 0.155 ± 0.005 | 0.32 ± 0.05 | 0.371 ± 0.016 | 0.57 ± 0.10 | 0.644 ± 0.033 | 0.78 ± 0.14 |
| *Tr*Cel7A | 0.056 ± 0.004 | 0.16 ± 0.06 | 0.138 ± 0.006 | 0.32 ± 0.06 | 0.250 ± 0.006 | 0.52 ± 0.05 | 0.307 ± 0.010 | 0.60 ± 0.07 |
| *Pc*Cel7D | 0.098 ± 0.003 | 0.13 ± 0.02 | 0.275 ± 0.007 | 0.21 ± 0.02 | 0.745 ± 0.016 | 0.45 ± 0.04 | 1.673 ± 0.030 | 0.92 ± 0.06 |
| *Tr*Cel7A$_{CD}$ | 0.084 ± 0.002 | 0.18 ± 0.02 | 0.226 ± 0.006 | 0.59 ± 0.06 | 0.551 ± 0.018 | 1.60 ± 0.15 | 0.641 ± 0.037 | 2.07 ± 0.33 |
| *Tr*Cel6A | 0.813 ± 0.027 | 3.30 ± 0.25 | 1.659 ± 0.104 | 6.00 ± 0.71 | 2.513 ± 0.199 | 6.33 ± 0.94 | 3.611 ± 0.295 | 6.94 ± 1.03 |
| *Tr*Cel7A$_{W38A}$ | 0.221 ± 0.007 | 0.83 ± 0.09 | 0.302 ± 0.013 | 1.10 ± 0.16 | 0.303 ± 0.015 | 0.75 ± 0.13 | 0.341 ± 0.027 | 1.00 ± 0.27 |
| *Tr*Cel6A$_{CcCBM}$ | 1.030 ± 0.026 | 5.35 ± 0.26 | 1.616 ± 0.065 | 6.59 ± 0.49 | 2.178 ± 0.136 | 7.19 ± 0.80 | 2.367 ± 0.051 | 5.58 ± 0.24 |
| *Tr*Cel5A | 1.019 ± 0.016 | 0.57 ± 0.03 | 2.174 ± 0.043 | 0.92 ± 0.06 | 4.355 ± 0.059 | 1.65 ± 0.06 | 7.074 ± 0.162 | 2.22 ± 0.13 |
| *Hi*Cel45A | 1.742 ± 0.094 | 0.99 ± 0.18 | 3.875 ± 0.124 | 2.34 ± 0.19 | 8.373 ± 0.408 | 4.94 ± 0.49 | 12.068 ± 0.302 | 6.02 ± 0.29 |
| *Tr*Cel12A | 0.847 ± 0.054 | 2.48 ± 0.41 | 0.737 ± 0.046 | 2.61 ± 0.41 | 0.851 ± 0.072 | 1.72 ± 0.42 | 0.563 ± 0.041 | 2.15 ± 0.42 |
| *Tr*Cel7B | 5.690 ± 0.202 | 3.66 ± 0.29 | 8.016 ± 0.174 | 3.98 ± 0.19 | 12.592 ± 0.328 | 5.24 ± 0.27 | 18.526 ± 0.503 | 7.20 ± 0.35 |
| *Rp*Cel45A | 1.249 ± 0.075 | 1.92 ± 0.32 | 1.706 ± 0.120 | 3.18 ± 0.53 | 2.475 ± 0.094 | 4.61 ± 0.37 | 2.090 ± 0.071 | 4.52 ± 0.32 |

N/A, parameter unable to determine accurately due to very low affinity.
*Enzyme not included in LFERs.
#Enzyme not included in LFERs due to unusual increase in affinity with temperature increase.

#### Linear free-energy relationships (LFERs) parameters

**Table S2.** Slope $\phi$, intercept $\beta$, and $R^2$ for cellulase LFERs at different substrates and temperatures.

| Substrate | $\phi$ | | | | |
|---|---|---|---|---|---|
| | ***20 °C*** | ***30 °C*** | ***40 °C*** | ***50 °C*** | ***Average*** |
| ***BMCC*** | 0.26 ± 0.08 | 0.28 ± 0.07 | 0.32 ± 0.08 | 0.26 ± 0.07 | 0.28 ± 0.03 |
| ***BC*** | 0.60 ± 0.09 | 0.52 ± 0.09 | 0.59 ± 0.10 | 0.64 ± 0.10 | 0.59 ± 0.04 |
| ***Avicel*** | 0.73 ± 0.08 | 0.76 ± 0.09 | 0.75 ± 0.11 | 0.73 ± 0.15 | 0.74 ± 0.01 |
| ***PCS*** | 0.96 ± 0.12 | 0.98 ± 0.12 | 0.93 ± 0.12 | 0.87 ± 0.13 | 0.93 ± 0.04 |
| ***RAC*** | 0.95 ± 0.20 | 0.88 ± 0.22 | 0.99 ± 0.24 | 1.16 ± 0.30 | 1.00 ± 0.10 |
| **Substrate** | $\beta$ | | | | |
| | ***20 °C*** | ***30 °C*** | ***40 °C*** | ***50 °C*** | |
| ***BMCC*** | -1.70 ± 0.12 | -1.18 ± 0.13 | -0.56 ± 0.16 | -0.16 ± 0.16 | |
| ***BC*** | -1.74 ± 0.11 | -1.40 ± 0.12 | -1.08 ± 0.18 | -0.91 ± 0.21 | |
| ***Avicel*** | -3.49 ± 0.27 | -3.31 ± 0.34 | -2.89 ± 0.44 | -2.52 ± 0.59 | |
| ***PCS*** | -2.72 ± 0.27 | -2.65 ± 0.30 | -2.32 ± 0.37 | -1.97 ± 0.40 | |
| ***RAC*** | -0.66 ± 0.25 | -0.47 ± 0.26 | -0.30 ± 0.29 | -0.42 ± 0.37 | |
| **Substrate** | $R^2$ | | | | |
| | ***20 °C*** | ***30 °C*** | ***40 °C*** | ***50 °C*** | ***Average*** |
| ***BMCC*** | 0.60 | 0.70 | 0.75 | 0.71 | 0.69 ± 0.06 |
| ***BC*** | 0.82 | 0.77 | 0.79 | 0.82 | 0.80 ± 0.02 |
| ***Avicel*** | 0.91 | 0.89 | 0.84 | 0.72 | 0.84 ± 0.07 |
| ***PCS*** | 0.86 | 0.87 | 0.85 | 0.83 | 0.85 ± 0.01 |
| ***RAC*** | 0.70 | 0.61 | 0.63 | 0.59 | 0.63 ± 0.04 |

#### Calorimetric measurements

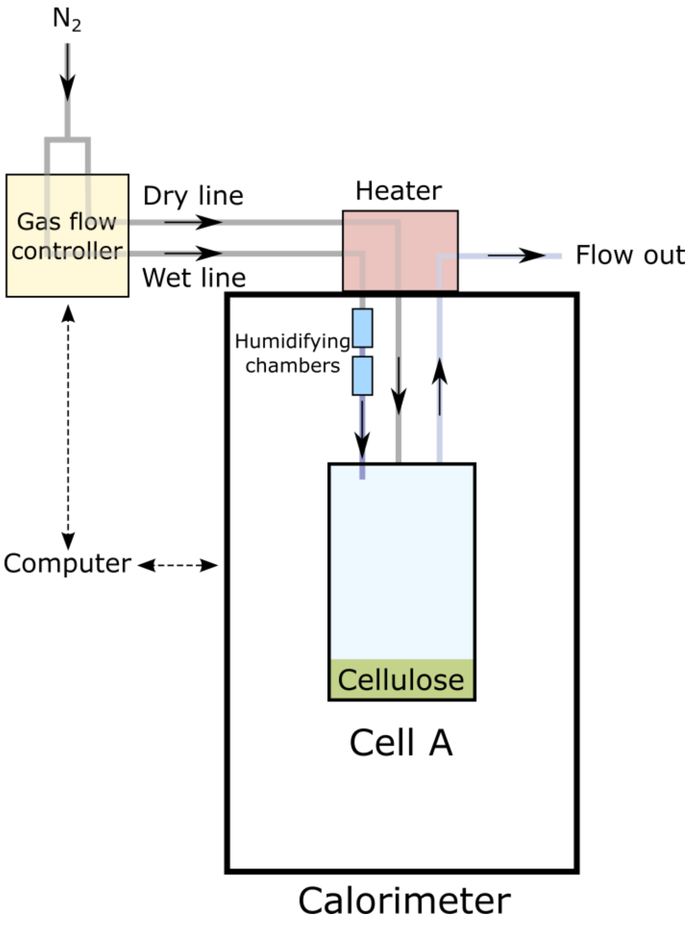


**Figure S1.** Schematic of the calorimetric setup.

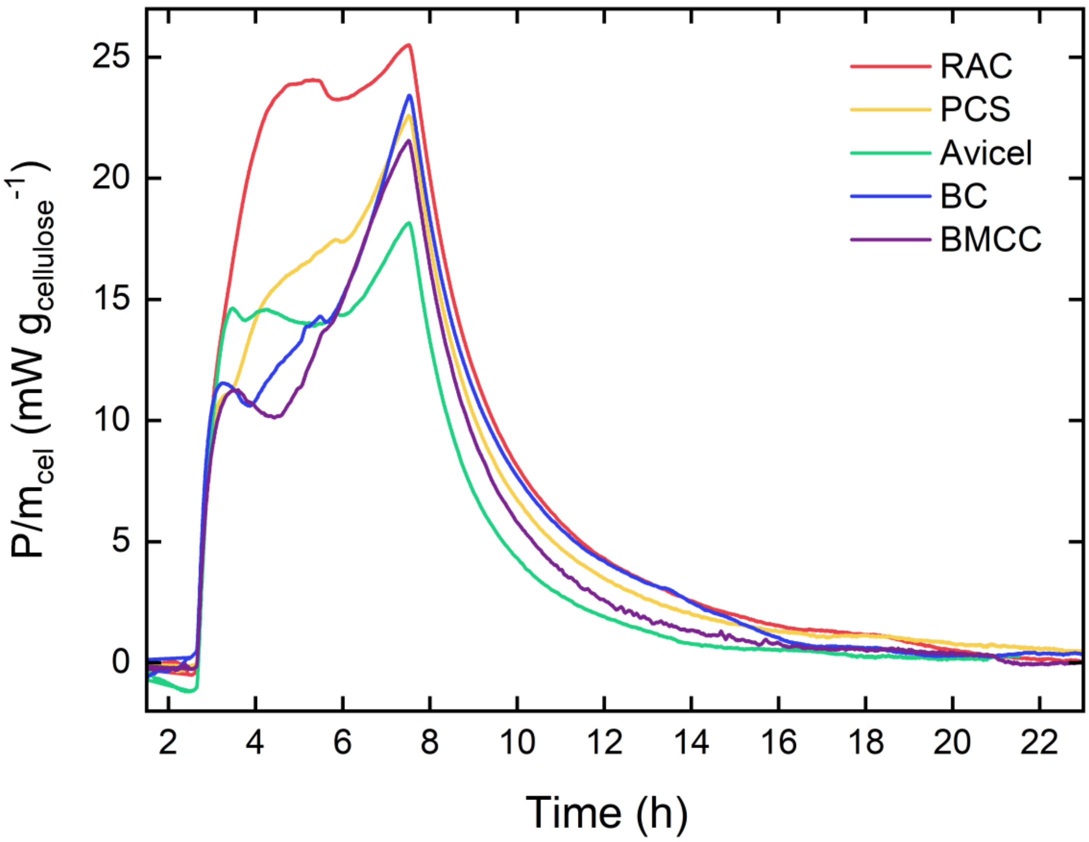


**Figure S2.** Example plots of RH perfusion calorimetry datasets from cell A of the cellulosic substrates investigated here. Blanks were subtracted from the raw data. The graph illustrates the heat flow P per cellulose dry mass $m_{cel}$ as a function of time. Dry gas (0% RH) was provided until t = 2.5 h, and then RH was raised to 96% at a rate of 19.2 %/h. The signal from t = 7.5 h to t = 23 h reflects gradual equilibration of the sorption process at RH = 96%.

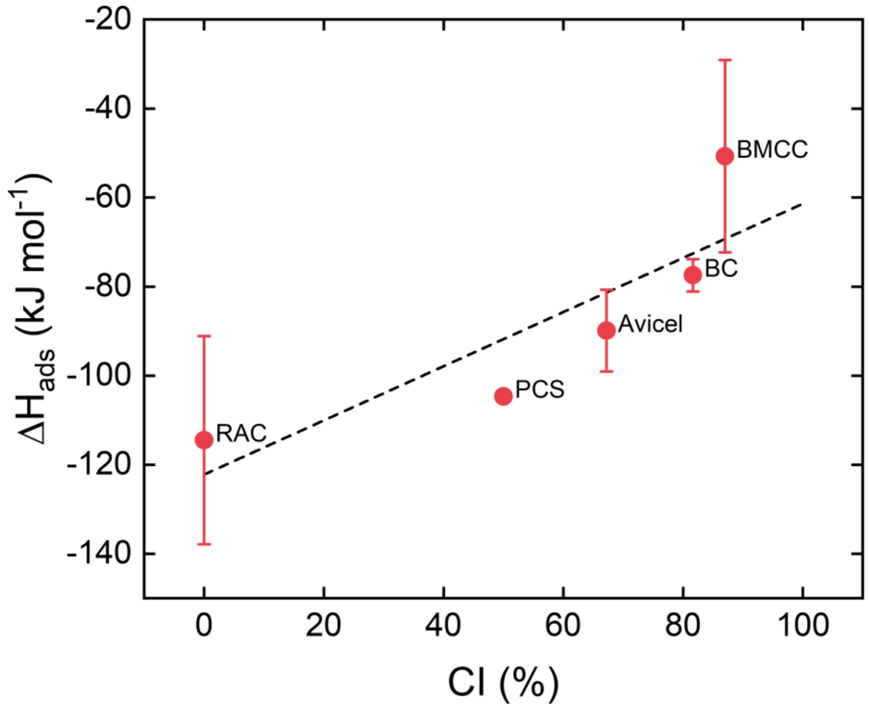


**Figure S3.** Plot of enthalpy of water adsorption $\Delta H_{ads}$ versus literature values of crystallinity index (CI) for the different celluloses investigated. The black dashed line represents the linear regression of this dataset. Error bars are standard deviations from tri- or quadruplicates, except for PCS, which is based on a single measurement.

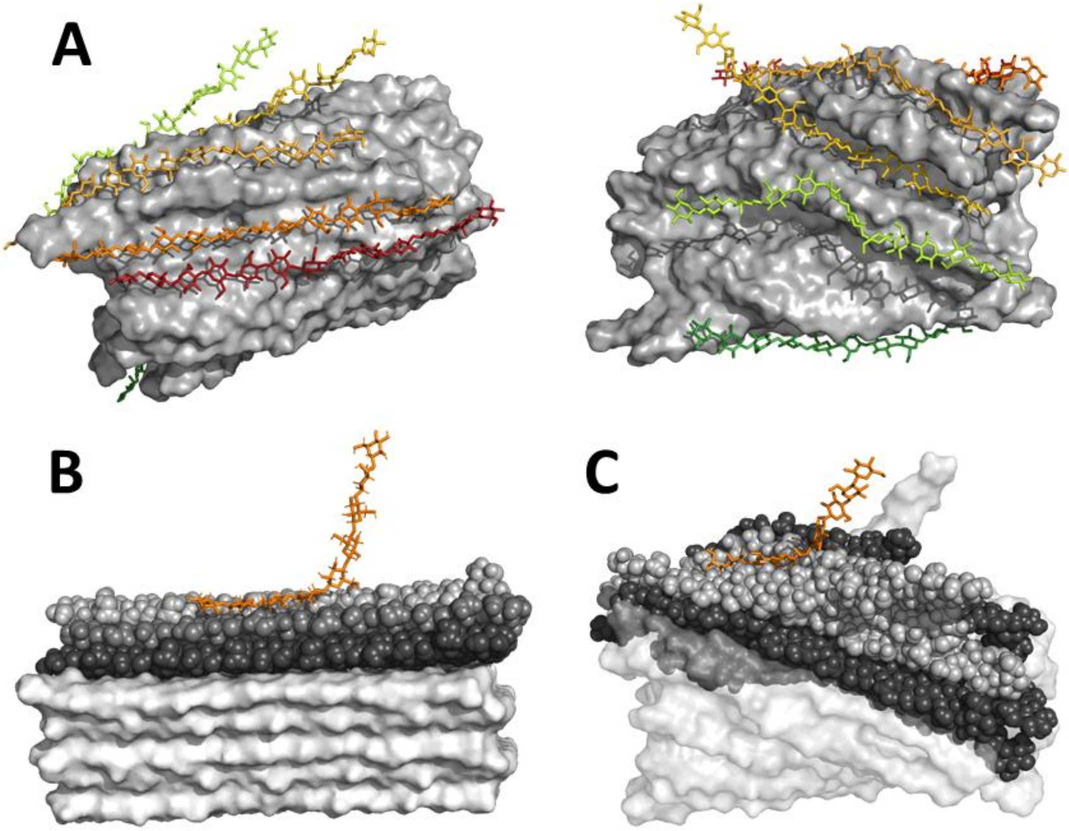


**Figure S4.** A) Part of an amorphous fibril from Vermaas *et. al* [1] from two sides with the different, individually investigated amorphous chains highlighted in color. B) Frame from the steered simulation of the crystalline substrate. The pulled nonaose is displayed as orange sticks, the upper 3 layers are shown as spheres. The third layer (black) was kept restraint. The remaining crystal fibril layers are rendered as a surface model for context, but were not part of the simulation. C) Frame from the steered simulation of one of the amorphous chains. The investigated nonaose is displayed as orange sticks, the first shell of surrounding chains are show as grey spheres, the second, restraint shell as black ones. The remaining amorphous fibril layers are rendered as surface model for context, but were not part of the simulation.

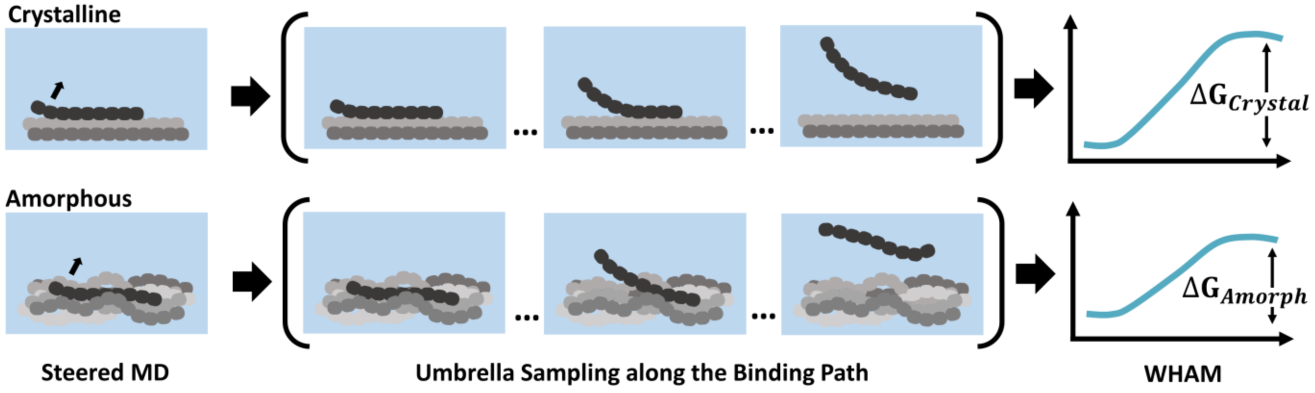


**Figure S5.** Schematic workflow to assess the detachment energies via umbrella sampling along the binding path with weighted histogram-analysis method (WHAM).[2] The chain of interest is slowly pulled from the initial structure to generate starting structures for the subsequent array of umbrella sampling simulations. The umbrella sampling simulations are then analyzed with WHAM to get an estimate of the free energy of ligand release.
